# Assessing genomic admixture between cryptic *Plutella* moth species following secondary contact

**DOI:** 10.1101/405498

**Authors:** Christopher M. Ward, Simon W. Baxter

## Abstract

Cryptic species are genetically distinct taxa without obvious variation in morphology and are occasionally discovered using molecular or sequence datasets of populations previously thought to be a single species. The world-wide Brassica pest, *Plutella xylostella* (diamondback moth), has been a problematic insect in Australia since 1882, yet a morphologically cryptic species with apparent endemism (*P. australiana*) was only recognized in 2013. *Plutella xylostella* and *P. australiana* are able to hybridize under laboratory conditions, and it was unknown whether introgression of adaptive traits could occur in the field to improve fitness and potentially increase pressure on agriculture. Phylogenetic reconstruction of 29 nuclear genomes confirmed *P. xylostella* and *P. australiana* are divergent, and molecular dating with 13 mitochondrial genes estimated a common *Plutella* ancestor 1.96±0.175 MYA. Sympatric Australian populations and allopatric Hawaiian *P. xylostella* populations were used to test whether neutral or adaptive introgression had occurred between the two Australian species. We used three approaches to test for genomic admixture in empirical and simulated datasets including i) the f3 statistic at the level of the population, ii) pairwise comparisons of Nei’s absolute genetic divergence (*d*_*XY*_) between populations and iii) changes in phylogenetic branch lengths between individuals across 50 kb genomic windows. These complementary approaches all supported reproductive isolation of the *Plutella* species in Australia, despite their ability to hybridize. Finally, we highlight the most divergent genomic regions between the two cryptic *Plutella* species and find they contain genes involved with processes including digestion, detoxification and DNA binding.

## Introduction

Cryptic species lack conspicuous variation in visible traits, yet can show high levels of ecological, behavioral and genetic divergence, particularly when they arise in allopatry (Bickford, et al. 2007; Pfenninger and Schwenk 2007; Stuart, et al. 2006). Morphological resemblance of two or more distinct species can occur when environmental pressures maintain phenotypes or cause convergence, and through introgression of traits by interspecies hybridization (Bickford, et al. 2007). Consequently, cryptic species are often overlooked, leading to both underestimates of species richness and overestimates of their geographic range (Stuart, et al. 2006; Voda, et al. 2015).

Reproductive barriers can maintain boundaries between sympatric congeneric animal species (cryptic or non-cryptic) using a range of isolating mechanisms such as olfaction, pheromone cues and mating calls (Andersson, et al. 2007; Jones and Hamilton 1998), host plant preference or mating timing (Hänniger, et al. 2017), and endosymbiont infection (Bordenstein, et al. 2001; Shoemaker, et al. 1999). Although these factors can impose reproductive isolation barriers and restrict hybridization, assortative mating does not always occur (Mallet, et al. 2007). Interspecific hybridization of two species within the same genera has been found to occur at similar rates across the animal kingdom, after taxonomic groups are adjusted for species richness (Schwenk, et al. 2008). While hybridization between related species has been well documented, the process of distinguishing between adaptive introgression and regions of historic population structure has been challenging (Martin, et al. 2015).

Closely related allopatric or sympatric species without gene flow should exhibit genetic divergence across the genome, whereas species with gene flow should show lower levels of divergence across broad regions relative to the frequency of interbreeding and how recently it occurred. Detecting hybridization is possible through the use of informal statistical tests on genetic variation, including principle component analysis (Patterson, et al. 2006) and Bayesian STRUCTURE model analysis (Pritchard, et al. 2000). While these tests can provide results indicative of admixture, they cannot distinguish between introgression, inter-lineage sorting or homoplasic genetic drift. Patterson, et al. (2012) formalized statistical approaches to estimate admixture based on allele frequencies across multiple populations, namely the f3 and f4 statistics (*D-*statistic), which assess the likelihood of hybridization. The f4 statistic has identified introgression between sympatric *Heliconius* butterfly species (Martin, et al. 2013; Zhang, et al. 2016) and hominids (Patterson, et al. 2012), as allele frequencies across these genomes did not always agree with the expected species tree, or neutral drift.

Hybridization and introgression of genetic variation from a donor species into a recipient can have adaptive advantages. The transfer of advantageous pre-adapted alleles from one species into another removes the reliance of new traits arising though mutation in the recipient. Examples include the transfer of rodenticide resistance between mice (Song, et al. 2011), coat colour alleles among jackrabbits and hares (Jones, et al. 2018), aposematic wing patterns in *Heliconius* butterflies (Mavárez, et al. 2006; Pardo-Diaz, et al. 2012; Wallbank, et al. 2016) and insecticide resistance genes in *Anopholes* mosquitoes (Lee, et al. 2013; Norris, et al. 2015).

The diamondback moth, *Plutella xylostella* (L.) (Lepidoptera: Plutellidae), is the most destructive pest of Brassicaceous agricultural crops, including broccoli, cabbage and canola (Furlong, et al. 2013). They are able to cause *en masse* defoliation, malformed and improper plant growth (Zalucki, et al. 2012), and often develop resistance to insecticides making pest control an ongoing challenge. *Plutella xylostella* were first documented in Australia in the 1880’s (Tyron 1889), yet an endemic and phenotypically cryptic species, *P. australiana* (Landry and Hebert), was only recently identified through high divergence of mitochondrial COI barcode sequences (8.6%) and morphologically distinct genitalia (Landry and Hebert 2013). The discovery was surprising, as *P. australiana* was not detected in previous molecular studies of *P. xylostella* yet, is dispersed across eastern Australia (Delgado and Cook 2009; Endersby, et al. 2006).

Insecticide susceptibility appears to limit *P. australiana’s* pest potential among cultivated brassica crops, however, introgression of insecticide resistance loci from *P. xylostella* could have serious consequences for agriculture. *Plutella xylostella* and *P. australiana* can hybridize in experimental laboratory crosses, despite their contrasting infection rates of endosymbiotic *Wolbachia* (Perry, et al. 2018; Ward and Baxter 2017), which are known to cause reproductive incompatibility in some cases (Duplouy, et al. 2013; Sasaki and Ishikawa 2000). *Wolbachia* infection is fixed among *P. australiana* yet extremely low in Australian *P. xylostella* (1.5%). Although the strength of reproductive barriers in the field is unknown, limited numbers of SNP markers widely dispersed across the nuclear genome previously identified genetic structure between sympatric populations of *P. xylostella* and *P. australiana* (Perry, et al. 2018). Due to *P. australiana’*s apparent endemism and the relatively recent invasion of *P. xylostella* into Australia, we assessed the capacity for sympatric Australian *Plutella* species to exchange beneficial traits through disassortative mating and introgression in the field through analyzing whole genomes.

## Materials and Methods

### Specimen collection and Genome Sequencing

*Plutella xylostella* and *P. australiana* were collected from canola (*Brassica napus*) fields using light traps at Cook, Australian Capital Territory (ACT), (−35.262, 149.058) in October 2014 and from direct larval sampling at Ginninderra Farm, ACT, (−35.187, 149.053) in December 2015. Larvae from Calca, South Australia, (SA) (−33.049, 134.373) and Bairds Bay, (SA) (−33.023, 134.279) were collected in June 2014 from mixed stands of sand rocket (*Diplotaxis tenuifolia*) and wall rocket (*D. muralis*). Larval collections were reared through to pupation then frozen, to eliminate samples infected with parasitoids. A single *P. australiana* moth was also collected from Richmond, New South Wales (−33.597, 15.740) using a light trap. Large populations of *P. xylostella* larvae were also collected from *Brassica* vegetable farms on three Hawaiian Islands in August 2013, including Kunia, Oahu (21.465, −158.064), Kula, Maui (20.791, −156.337) and Waimea on Hawaii Island (20.028, −155.636), and reared for one generation. Genomic DNA purification was performed using phenol extractions, treated with RNaseA, precipitated with ethanol and resuspended in TE buffer (10 mM Tris, 0.1 mM EDTA). Species identification was performed using a PCR-RFLP diagnostic assay of the mitochondrial COI gene (Perry, et al. 2018). Genome sequencing was performed using the Illumina HiSeq2500 or NextSeq platforms at the Australian Genome Research Facility and the Australian Cancer Research Facility.

### Processing genome sequence data

Summary statistics of Illumina sequence reads were generated with FastQC (Andrews 2010) and visualized using the R package ngsReports (Ward, et al. 2018). Trimmomatic v 0.32 (Bolger, et al. 2014) was used with the parameters [TRAILING:15 SLIDINGWINDOW:4:15] to trim adapter, quality filter and retain paired reads. The *P. xylostella* reference genome (You, et al. 2013) was downloaded from NCBI (GCA_000330985.1). Stampy v1.0.21 (Lunter and Goodson 2011) was used to align the paired reads to the reference with the parameters [*--gatkcigarworkaround, -- substitutionrate=0.01*] which produced Sequence Alignment/Map (SAM) files that were converted to binary format (BAM) and indexed then sorted using SAMtools v1.2 (Li, et al. 2009). PCR and optical duplicates were removed using Picard Tools v1.61 (http://broadinstitute.github.io/picard/). BAM summary statistics including average read depth per site called, coverage of the genome, percent missing data, total number of reads and read quality were generated using SAMtools v1.2.

### Genotype Variant Calling

Variant calling was performed using the Genome Analysis ToolKit (GATK) v3.3 (DePristo, et al. 2011). GATK:HaplotypeCaller was used to generate gVCF records, containing variant and invariant sites across the genome, on a per sample basis. The HaplotypeCaller parameter heterozygosity (likelihood of a site being non-reference) for each species was estimated by SAMtools v1.2, indicating *P. xylostella* from Hawaii was most similar to the reference genome (heterozygosity: *P. australiana* = 0.0497; *P. xylostella* Australia = 0.0348; *P. xylostella* Hawaii = 0.0272). Individual gVCF records were combined using GATK:Genotype GVCF and filtered using BCFtools (Li, et al. 2009) to a minimum individual depth greater than 5 reads per base with no greater than 40% of samples missing genotypes at any one site.

Nei’s mean intra-population nucleotide diversity, π, (Nei and Li 1979) was calculated using egglib (De Mita and Siol 2012). The mean and standard error in π and jackknifing was performed using the R package bootstrap (Canty and Ripley 2017). Pairwise F_ST_ and Tajima’s D was calculated across 50 kb windows using VCFtools (Danecek, et al. 2011) and minimum distances between populations (km’s) determined with http://www.movable-type.co.uk/scripts/latlong.html.

### Phylogenetic reconstruction of *Plutella* mitochondrial and nuclear genomes

All quality filtered variant and invariant sites called against the mitochondrial reference genome (GenBank KM023645) were extracted using BCFtools and converted to a FASTA alignment using the R programming language. Maximum likelihood phylogenetic inference using the nuclear genome consensus, heterozygous sites were replaced with IUPAC ambiguity codes, alignment was performed with exaML (Kozlov, et al. 2015) with GTR+GAMMA bootstrap resampling (n=100; GTR+GAMMA) was then carried out using RAxML v8.2.4 to provide node confidence. The phylogeny was then rooted using the mid-point method in FigTree (v1.4.3, http://tree.bio.ed.ac.uk/software/figtree)

### *De novo* assembly of mitochondrial genomes and *Plutella* split time estimates

*De novo* assembly of *Plutella* mitochondrial genomes was performed using NOVOPlasty v2.6.3 (Dierckxsens, et al. 2017). A sequence read that mapped to the *P. xylostella* mitochondrial COI gene was used as the seed to initiate assembly. Genomes circularized by NOVOPlasty were then annotated through homology to the *P. xylostella* mitochondrial reference gene annotation (GenBank KM023645) with Geneious v10.0.6. Potential misassemblies were investigated by mapping individual raw reads to the appropriate *de novo* assembly on a per sample basis using BWA-MEM (Li 2013). Mapped reads were then used as fragments in Pilon (Walker, et al. 2014) to correct the assembly. The sample with the greatest total length (15962 bp), *Paus ACT14.1*, was used to produce a reference for the mitochondrial genome of *Plutella australiana* (Genbank accession MG787473.1).

The mitochondrial split time between *P. xylostella* and *P. australiana* was estimated using 13 mitochondrial protein coding genes extracted from 20 *Plutella* samples with circularized genomes plus *Prays oleae* (accession no. NC_025948.1) and *Leucoptera malifoliella* (accession no. JN790955.1). Nucleotide alignments were made for each gene using MAFFT (Katoh, et al. 2002), substitution models were determined using JModelTest2 (Darriba, et al. 2012) and alignments were then imported into BEAUTi (Drummond, et al. 2012). We set the clock model to strict with 0.0177 substitutions My^-1^ according to Papadopoulou, et al. (2010). Substitution models were unlinked to allow each sequence to coalesce independently with the Yule speciation model. MCMC sampling was carried out over 1000000 trees sampling every 1000 using BEAST2 v 2.4.7 (Bouckaert, et al. 2014). Sampled trees from the chain were checked using Tracer v 1.6 (Rambaut, et al. 2014) to determine burn in. Densitree was then used to superimpose MCMC trees to determine the internal node height ranges.

### Data simulation

Coalescent local trees with a total chromosomal length of 25 Mb were simulated for 24 individuals, including eight samples from an outgroup (O) and two ingroups (I_1_ and I_2_) using the Markovian Coalescent Simulator, MaCS (Chen, et al. 2009). A coalescent model for the most recent common ancestor of I_1_ and I_2_ was set to 0.4 × 4N generations ago and the root to 1.5 × 4N generations ago, providing the topology ((I_1_, I_2_), O). Simulated divergence was determined using mean *d*_*XY*_ values from *Plutella* samples (see Figure 4). Two approaches were used to simulate introgression events from I_2_ to O or from O to I_2_. First, Introgression was simulated as a single *en masse* admixture event at 0.01 × 4N generations ago with admixture frequencies (*f*) of *f =* 0, 0.05, 0.1, 0.2 and 0.3. Second, introgression was simulated over five distinct breakdowns in assortative mating (0.01, 0.008, 0.006, 0.004 and 0.002 x 4N generations ago) and *f =* 0, 0.05, 0.1, 0.2 and 0.3. Each simulation was carried out with a constant population recombination rate (4Nr) of 0.001. Sequences were generated from the coalescent trees using SeqGen (Rambaut and Grassly 1997) with the Hasegawa-Kishino-Yano substitution model (Hasegawa, et al. 1985) and a branch scaling factor of 0.01.

### Admixture

#### The F3-statistic

A formal test for admixture was calculated using the three population test, the f-statistic (f3) (Patterson, et al. 2012; Reich, et al. 2009). Three possible combinations of tip structures were assessed with the ingroups and outgroup namely f3(I_1_, I_2_; O), f3(I_1_, O; I_2_), f3(I_2_, O; I_1_,). Cases without introgression are expected to return positive f3 values while negative values indicate introgression has occurred from a donor to a recipient population, forming an intermediate ancestor of both source populations. Block jack-knife F3 estimation was carried out using PopStats (https://github.com/pontussk/popstats).

#### Absolute divergence (d_XY_)

Nei’s absolute divergence, *d*_XY_, was used to calculate the mean number of nucleotide differences between two populations across non-overlapping 50kb windows with *egglib_sliding_windows.py* (https://github.com/johnomics). Comparisons of *d*_*XY*_ were made first with simulated datasets, using the five admixture frequencies (*f* = 0, 0.05, 0.1, 0.2 and 0.3) between I_2_ and O, O and I_2_ and I_1_ and I_2_. The *d*_*XY*_ values were summarized by transforming them into density plots to visualize the distribution and frequency across the simulated genome. This provided expected *d*_*XY*_ patterns under a range of admixture frequencies. Average *d*_*XY*_ was then calculated between i) *P. australiana* (O) and Australian *P. xylostella* (I_2_) individuals, ii) *P. australiana* (O) and Hawaiian *P. xylostella* (I_1_) and iii) Hawaiian *P. xylostella* (I_1_) and Australian *P. xylostella* (I_2_). Histograms were plotted after setting the maximum value to 1 using the *geom_density* function in ggplot2 (Wickham 2009).

#### Tree-Tip Distance Proportions

Maximum likelihood phylogenetic reconstruction was performed with RAxML (Stamatakis 2014) using non-overlapping 50 kb genomic windows generated with the python script *genoToSeq.py* (https://github.com/simonhmartin). Phylogenies of empirical data each used four individuals, including one *P. xylostella* and one *P. australiana* individual from sympatric Australia populations (SA14, ACT14 or ACT15), and two *P. xylostella* individuals from Hawaii (HO13.1 and HH13.2). Each tree was then converted to a distance matrix using APE (Paradis, et al. 2004) and pairwise distances between tips were determined in the R programming language using the equation,

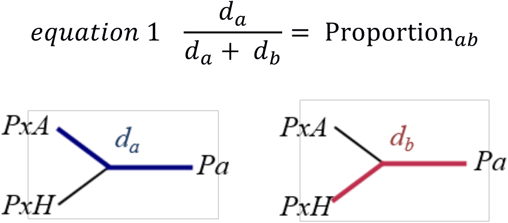

where *d*_*a*_ is the branch distance between the tree-tip of an Australian *P. xylostella* individual (*PxA*) and a *P. australiana* individual (*Pa*) and *d*_*b*_ is the averaged branch distance between two Hawaiian *P. xylostella* [*Pxyl HO13.1* and *Pxyl HH13.2*] and a *P. australiana* individual. Two individuals from Hawaii were used to reduce bias from this ingroup source. The *Proportion*_*ab*_ values for each 50 kb window were expected to be approximately 0.5 if the phylogeny was concordant with the species tree. Values much greater than 0.5 indicate genomic windows more similar between *P. australiana* and Hawaiian *P. xylostella*, while values much less than 0.5 indicate genomic windows more similar between *P. australiana* and Australian *P. xylostella* and are candidate admixed regions. Only genomic windows with more than 20% of sites genotyped were analyzed.

For comparison, simulated data from 24 individuals and five admixture frequencies (*f* = 0, 0.05, 0.1, 0.2 and 0.3) described above was divided into 50 kb windows (n = 500). Each 50 kb window was then subdivided into 64 separate alignments containing four simulated samples; the same two I_1_ individuals in each case, (reflecting the use of the same two *P. xylostella* samples from Hawaii in the empirical data) and non-redundant pairs of I_2_ and O individuals. Four tip unrooted maximum likelihood phylogenies were produced for each alignment using RAxML, then Proportion_*ab*_ calculated and plotted using a bin width of 0.05.

### Analysis of discordant tree-tip distances

After plotting tree-tip distance proportions, the tails of each distribution was investigated for symmetry by counting the number of 50 kb windows above or below each mean at three thresholds (mean ± 0.05, 0.10 and 0.15). Windows below the mean (mean – 0.15) were further investigated by calculating *d*_*XY*_ across 10 kb windows, sliding by 2 kb. These genomic regions indicate greater similarity between *P. australiana* and Australian *P. xylostella* than the average and *d*_*XY*_ plots were visually inspected for signs of introgression.

### Identification of divergent genomic windows between *P. australiana* and *P. xylostella*

Both F_ST_ *and d*_*XY*_ were calculated across aligned 50 kb genomic windows between all *P. xylostella* samples (from Australia plus Hawaii) and *P. australiana.* Annotated protein coding genes were extracted from the most divergent 1% of 50 kb windows for each statistic and BLAST against the DBM gene list available from DBM-DB (Tang, et al. 2014). To identify their molecular function, InterPro and UniProt annotations were obtained for each BLAST hit.

## RESULTS

### Alignment of *Plutella* species to the reference genome

The genomes of 29 *Plutella* samples were sequenced using short read Illumina platforms, including eight *P. xylostella* from Hawaii, eight *P. xylostella* from Australia and 13 *P. australiana*. Samples from Australia were classified into three populations based on collection location and year for analysis (ACT2014, ACT2015, SA2014). A single *P. australiana* individual from Richmond, NSW, was also sequenced (**Table S1**). Resequenced genomes were mapped to the ∼393 Mb *P. xylostella* reference genome (You, et al. 2013), but just 170 Mb of non-N bases were retained after stringent quality filtering. Sequence coverage across the 170 Mb alignment ranged from 9 to 25-fold per individual and approximately 70% of these sites were genotyped in *P. australiana* samples compared to ∼92% for Australian and Hawaiian populations of *P. xylostella* (**Table 1**).

**Table 1.** Summary of sequence coverage, percentage of sites genotyped and mean nucleotide diversity of *Plutella* populations (170 Mb)

| Population | Year collected | Species | Number of samples | Average coverage per site | Sites genotyped (%) | Mean nucleotide diversity | (±SD) |
| --- | --- | --- | --- | --- | --- | --- | --- |
| Australian Capital Territory (ACT) | 2014 | *P. xylostella* | 2 | 17 | 91.5 | 0.0151 | (0.0042) |
|  |  | *P. australiana* | 4 | 12.5 | 71.8 | 0.0170 | (0.0052) |
| Australian Capital Territory (ACT) | 2015 | *P. xylostella* | 2 | 18 | 92.2 | 0.0150 | (0.0045) |
|  |  | *P. australiana* | 4 | 13.5 | 72.3 | 0.0168 | (0.0048) |
| South Australia (SA) | 2014 | *P. xylostella* | 4 | 23.5 | 91.8 | 0.0157 | (0.0040) |
|  |  | *P. australiana* | 4 | 17.5 | 65.9 | 0.0174 | (0.0051) |
| Hawaii | 2013 | *P. xylostella* | 8 | 13.125 | 91.7 | 0.0200 | (0.0044) |

The highest levels of nucleotide diversity were observed within Hawaiian *P. xylostella* samples (**Table 1**). However, endemic *P. australiana* populations showed higher levels of nucleotide diversity than Australian *P. xylostella*, which may have undergone a population genetic bottleneck when colonization occurred. Mutation-drift equilibrium of these populations was determined using Tajima’s D (D_T_). *P. xylostella* collected from Australia were under equilibrium (D_T_ 95% CI = −0.6046375 to +0.9435148) whereas those collected from Hawaii showed largely negative values (D_T_ 95% CI = −1.88 to −0.039) which may be the result of a recent population size expansion or higher than expected abundance of rare alleles. The frequency of rare alleles in *P. australiana* was also common, although the D_T_ 95% confidence interval overlapped with zero (D_T_ 95% CI = −1.18 to + 0.42).

Pairwise comparisons between populations and species were then used to assess genetic structure with F_ST_. The three Australian *P. xylostella* populations showed no genetic structure between geographic location (SA vs. ACT) or year (2014 vs. 2015) (combined average of F_ST_ =0.003±0.003), as has been previously reported with microsatellite data (Endersby, et al. 2006). However, much higher levels of differentiation were observed when compared to Hawaiian *P. xylostella*, supporting the expectation of genetic isolation (average of F_ST_ =0.108±0.01). The average pairwise F_ST_ values were slightly lower between *P. australiana* and Hawaiian *P. xylostella* (F_ST_ =0.501±0.002) than *P. australiana* and Australian *P. xylostella* (F_ST_ =0.532±0.013) (**Table 2**).

**Table 2.**
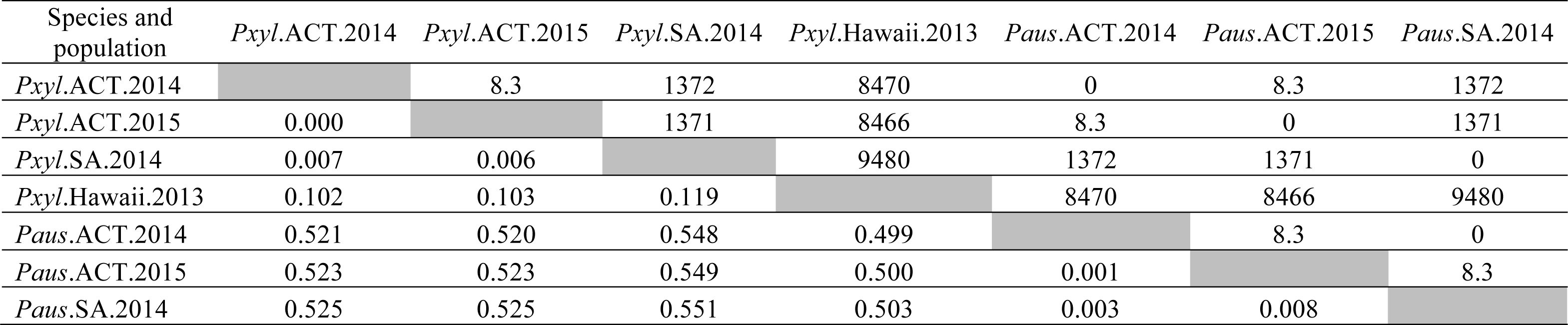
Matrix of the minimum distance between collection sites (km’s, above diagonal) and pairwise F_ST_ values of each *Plutella* population (below diagonal).

| Species and population | *Pxyl*.ACT.2014 | *Pxyl*.ACT.2015 | *Pxyl*.SA.2014 | *Pxyl*.Hawaii.2013 | *Paus*.ACT.2014 | *Paus*.ACT.2015 | *Paus*.SA.2014 |
| --- | --- | --- | --- | --- | --- | --- | --- |
| *Pxyl*.ACT.2014 |  | 8.3 | 1372 | 8470 | 0 | 8.3 | 1372 |
| *Pxyl*.ACT.2015 | 0.000 |  | 1371 | 8466 | 8.3 | 0 | 1371 |
| *Pxyl*.SA.2014 | 0.007 | 0.006 |  | 9480 | 1372 | 1371 | 0 |
| *Pxyl*.Hawaii.2013 | 0.102 | 0.103 | 0.119 |  | 8470 | 8466 | 9480 |
| *Paus*.ACT.2014 | 0.521 | 0.520 | 0.548 | 0.499 |  | 8.3 | 0 |
| *Paus*.ACT.2015 | 0.523 | 0.523 | 0.549 | 0.500 | 0.001 |  | 8.3 |
| *Paus*.SA.2014 | 0.525 | 0.525 | 0.551 | 0.503 | 0.003 | 0.008 |  |

### Phylogenetic inference of *Plutella* species

A maximum likelihood phylogeny using ∼170 Mb of the nuclear genome showed two clear *Plutella* species groups with deep divergence between species. *Plutella xylostella* from Hawaii and Australia formed reciprocally monophyletic sister clades with 100% bootstrap support while *P. australiana* genomes formed a single clade, although generally had lower levels of internal branch support (**Figure 1**). Branch distances were shorter between the internal nodes of *P. australiana* and Hawaiian *P. xylostella* than Australian *P. xylostella*, suggesting the two *P. xylostella* clades have diverged substantially since their most recent common ancestor.

**Fig. 1:**
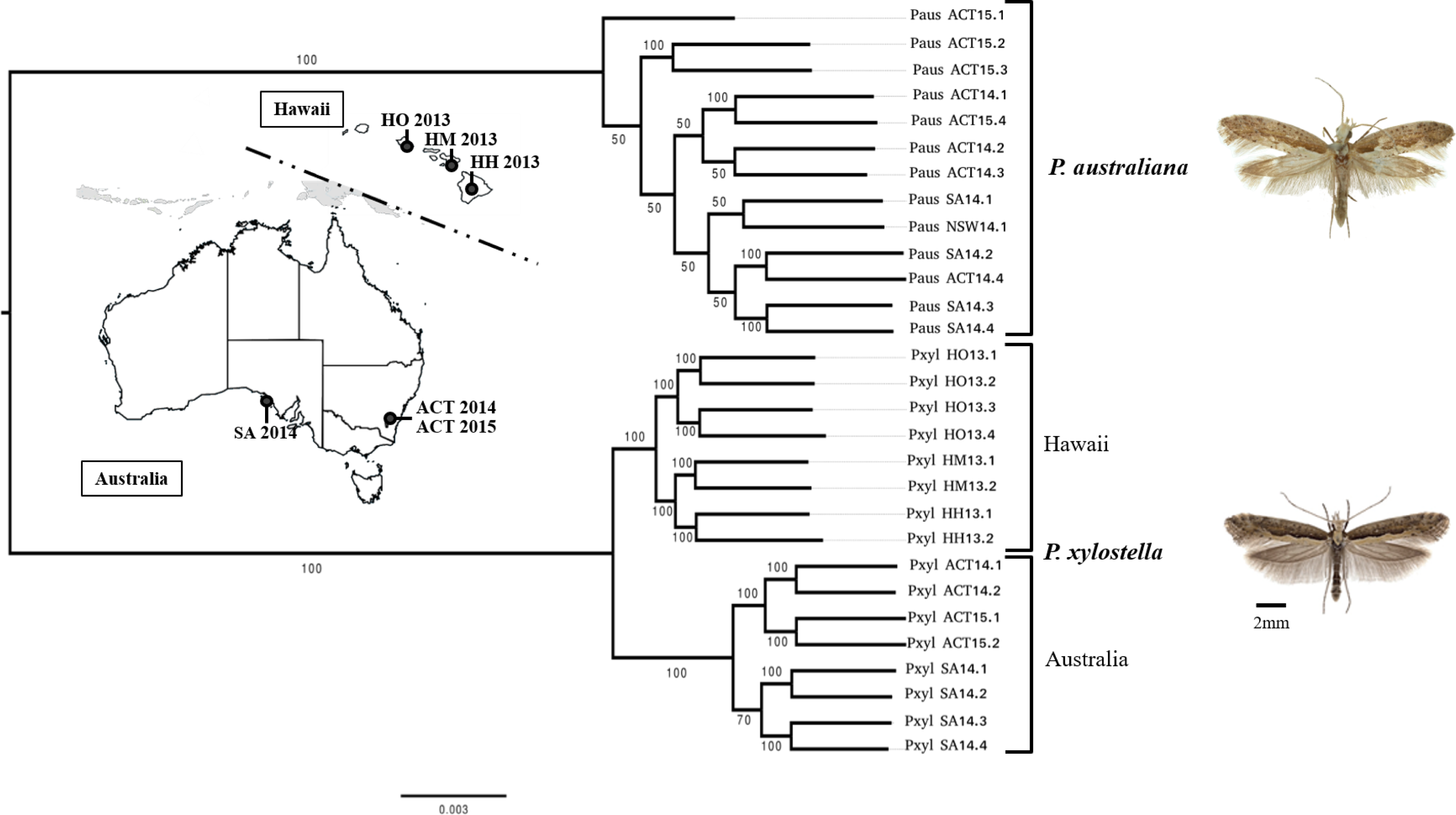
Maximum likelihood phylogeny of *P. xylostella* and *P. australiana* generated using a 170 Mb concatenated alignment of the nuclear genome. Bootstrap support (n=100) is shown at each node. The inner maps show population locations and year collected for samples from Australia (SA, South Australia; ACT, Australian Capital Territory) and Hawaii, USA (HO, Hawaii Oahu; HM, Hawaii Maui; HH, Hawaii, Hawaii Island). See Table 2 for distances between collection locations. Insect photographs were provided by Paul Hebert (*P. australiana*) and Jean-François Landry (*P. xylostella*).

### *P. australiana* mitochondrial genome and dating

We carried out *de novo* assembly and annotation of the *P. australiana* mitochondrial genome which has a total length of 15, 962 bp (GenBank accession MG787473.1) compared with 16, 014 bp of *P. xylostella* (Dai, et al. 2016b). Using sequence homology to the *P. xylostella* mitochondrial genome we annotated two rRNAs, 13 protein coding mitochondrial genes and 22 t-RNA, which showed a conserved gene order for Lepidopteran mitochondrial genomes (Dai, et al. 2016a). The nucleotide sequence of 13 protein coding mitochondrial genes from 22 *Plutella* samples were then used to estimate the mitochondrial split time between *P. xylostella* and *P. australiana* at 1.96 Mya (95% confidence interval ± 0.175 My **Figure 2**, **Table S2**). *Prays oleae* and *Leicoptera malifoliella* were used as the outgroups. The topology of the 13 mitochondrial genes used to date the split (**Table S3**) also supported two clear *Plutella* species groups with an average of 4.95% divergence. *Plutella xylostella* from Hawaii showed higher mitochondrial diversity than samples from Australia. Reduced mitochondrial diversity may have been caused by a founder effect when Australia was colonized (Perry, et al. 2018).

**Fig. 2.**
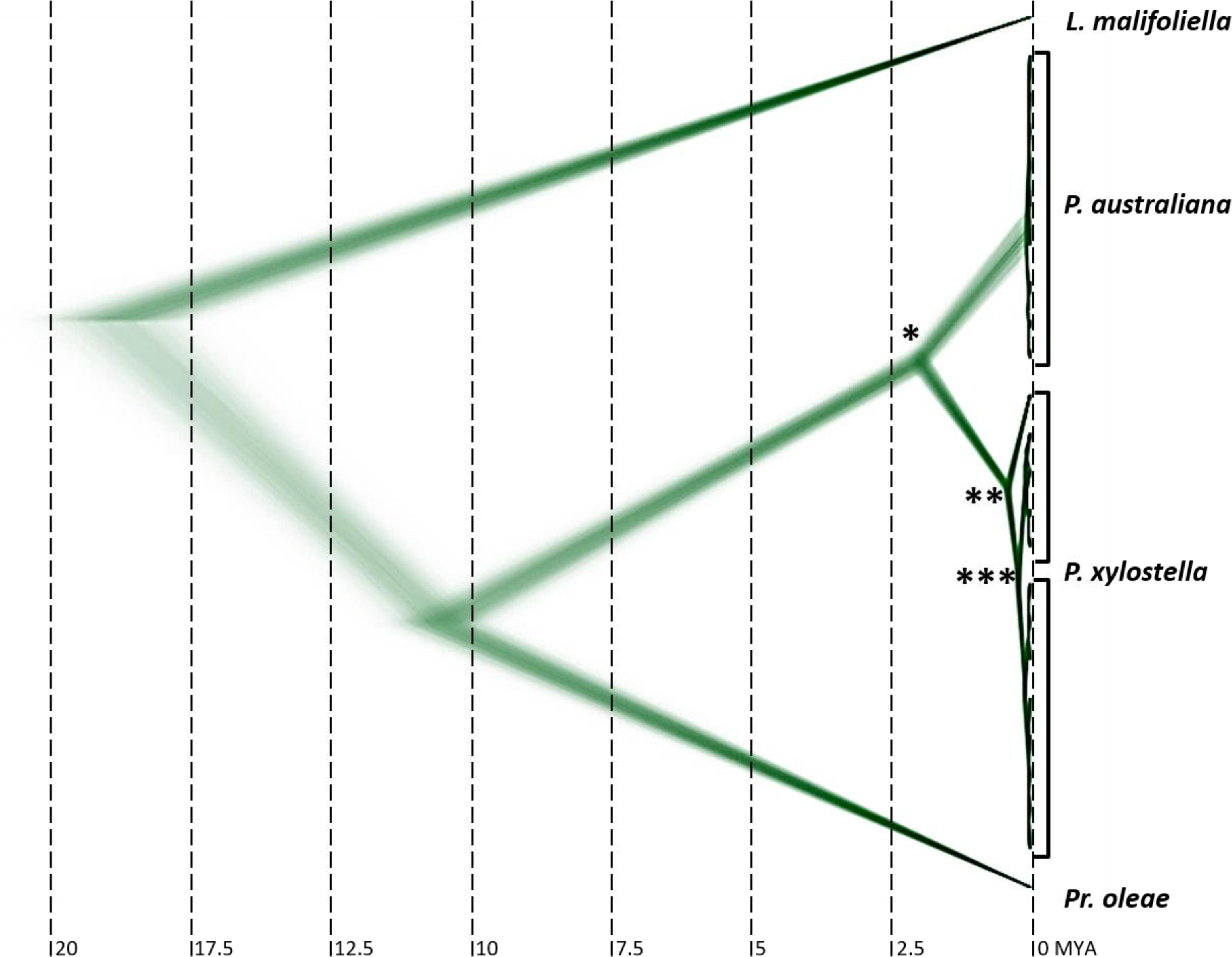
Superimposed MCMC trees of 13 protein coding mitochondrial genes used to estimate the split time of *P. xylostella* (n = 13) and *P. australiana* (n = 9) at 1.96 ± 0.175 Mya (*). The internal node of the *P. xylostella* clade was estimated at 0.37 ± 0.057 Mya (**). The split of Hawaiian and Australian *P. xylostella* haplotypes was estimated at 0.078 ± 0.024 Mya (***). *Prays oleae* (accession no. NC_025948.1) and *Leicoptera malifoliella* (accession no. JN790955.1) were used as outgroups.

### Assessing admixture between Australian *Plutella* species

#### The F3-statistic

A formal test for genomic admixture was calculated using the three-population f-statistic (f3), first with simulated datasets to assess the level of sensitivity we could reasonably achieve, and second with empirical data. Simulated introgression frequencies of *f=*0.0, 0.05, 0.1, 0.2 and 0.3 were applied from a donor to a recipient. Introgression from ingroup 2 (I_2_) into the outgroup (O) increased similarity between these groups, yet also reduced genetic differences between the outgroup and ingroup 1 (I_1_). Despite the outgroup becoming more similar to I_2_, O still contained a large proportion of divergent loci which tends to confound the f3 statistic making negativity difficult to achieve, even with high levels of introgression (Peter 2016). Consequently, this approach failed to indicate shared ancestry through a negative f3-statistic (**Figure 3A**). Next, introgression from O into I_2_ was simulated, to assess sensitivity of introgression from *P. australiana* into Australian *P. xylostella*. Negative values were detected for mixing frequencies of ≥20% (*f*=0.2), indicating high rates of recent hybridization are required to detect introgression using the f3-statistic (**Figure 3B**). Interestingly, spreading the total proportion of introgression to five equidistant time-points along the branch did not increase the detectability of admixture (**Figure S1**). This suggests the f3 is more dependent on the admixture frequency than the divergence between discordant and concordant regions.

**Fig. 3.**
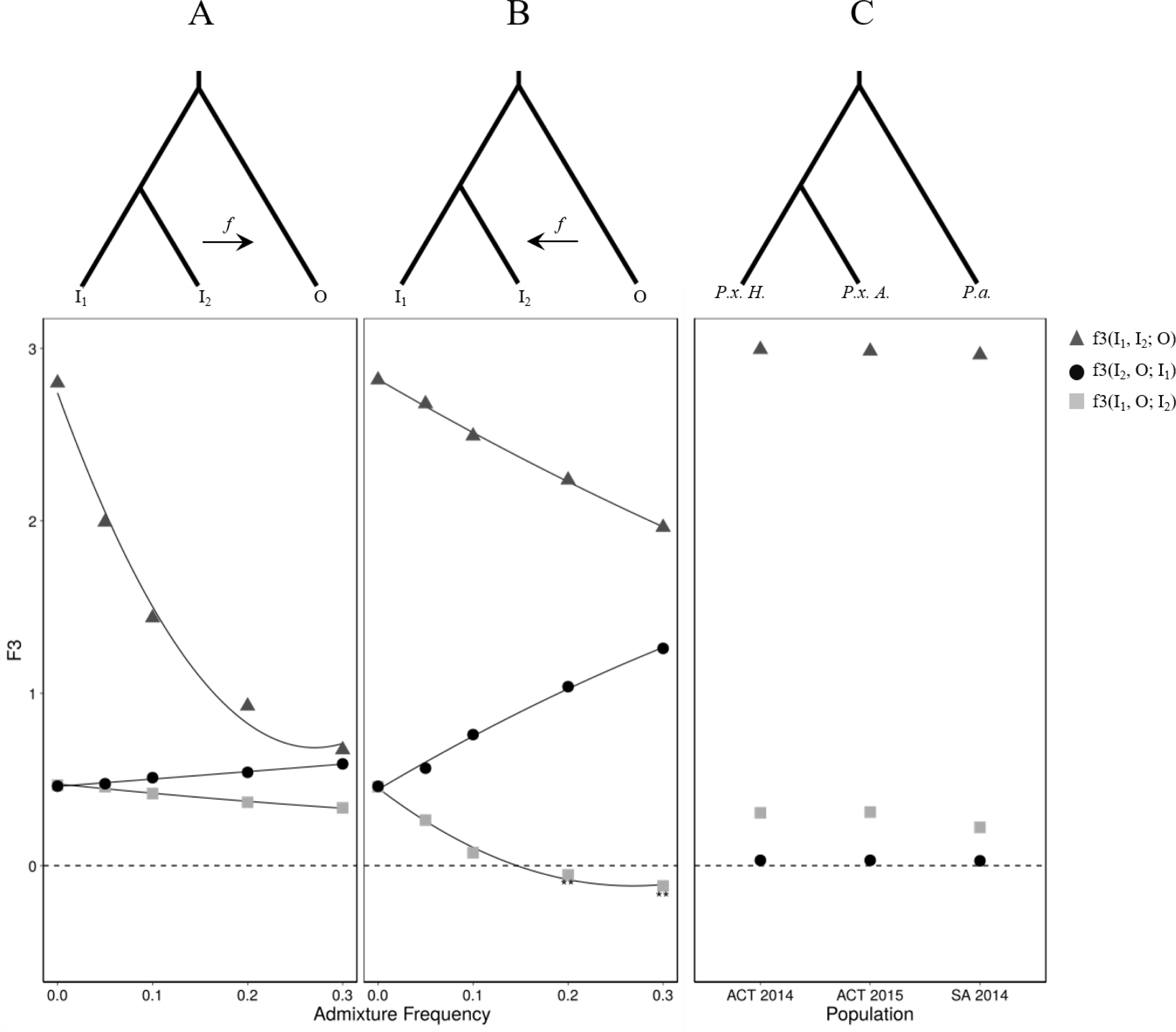
The three population f-statistic (f3). **A** Admixture from ingroup 2 (I_2_) to the outgroup (O) was simulated as a single event with frequencies of *f* = 0, 0.05, 0.1, 0.2 and 0.3. Evidence for hybridization and admixture could not be clearly detected in this direction, as shown by the grey boxes for f3(I_1_,O;I_2_), which did not reach negative values. For comparison, the f3-statistic for (I_2_,O;I_1_) and (I_1_,I_2_;O) were plotted with circles and triangles, respectively. **B** Simulated admixture from O into I_2_ did produced a significant f3 statistic at a mixing frequency >0.2, as indicated by the values below zero f3(I_1_,O;I_2_). **C** The f3-statistic was then applied to empirical data, testing for admixture in three possible scenarios between *P. xylostella* from Hawaii (*P.x. H,* I_1_), *P. xylostella* from Australia (*P.x. A,* I_2_) and *P. australiana* (*P.a.,* O). This suggests that, as no f3 values were below zero, if admixture was occurring it could not be detected using this method.

**Fig. 4.**
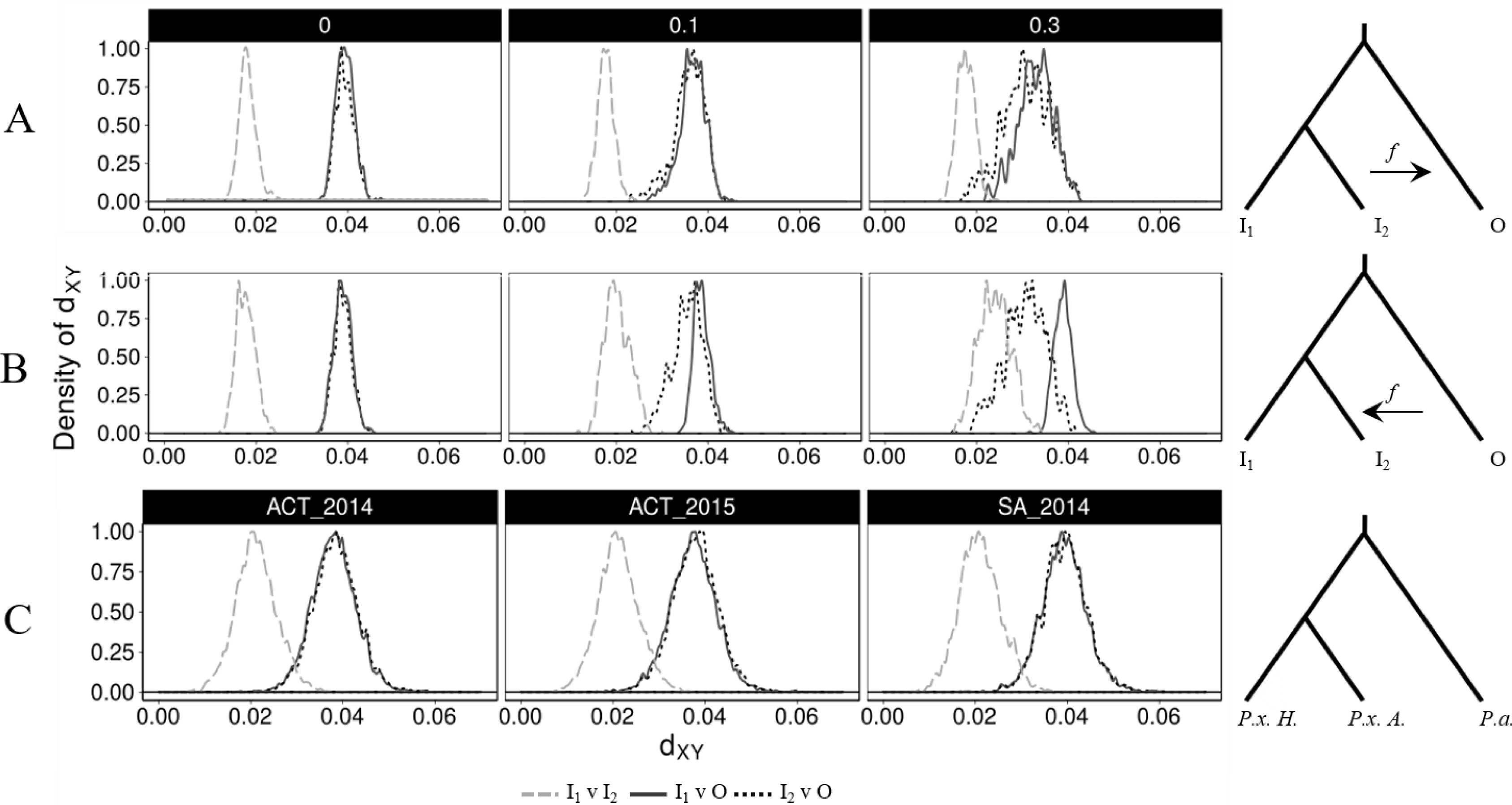
Absolute genetic divergence (*d*_*XY*_) between populations. Each plot is a histogram summarising pairwise comparisons of 50 kb windows across the genome, rescaled such that the maxima is 1. Simulated data uses mixing frequencies of *f*= 0.0, 0.1 and 0.3 (see **Figure S2** and **S3** for additional admixture frequencies) **A.** Simulated *d*_*XY*_ comparisons assessing of admixture from I_2_ into O or **B.** O into I_2_. In the abseence of hybridization (*f=*0) the simulated ingroups show the lowest levels of divergence (dashed line), while the distance between each ingroup and the outgroup are relatively similar (dotted and solid lines). Increasing levels of admixture (*f*=0.1, 0.3) alters histogram shape as I_2_ and O *d*_*XY*_ values become smaller. **C.** *d*_*XY*_ summaries between *P. australiana* (*P.a.* in the phylogeny schematic) and *P. xylostella* from Australia (*P.x. A*) or Hawaii (*P.x. H*) do not deviate. This indicates *d*_*XY*_ was not able to detect hybridization at the population level.

Applying the f3-statistic to empirical data failed to identify negativity in any tip order between Australian *P. xylostella* and *P. australiana* (**Figure 3C**). Results for the f3-statistic were lower when assessing introgression between Hawaiian *P. xylostella* and *P. australiana* than from between the two sympatric Australian species, consistent with the nuclear phylogeny showing Hawaiian samples are more similar to *P. australiana*. A lower f3(*Pxyl* Hawaii, *Pau.*; *Pxyl* Australia) value was estimated for SA 2014 than ACT 2014 and ACT 2015, which may be due to differences in nucleotide diversity between the Australian populations (**Table 1**), as f3 is decreased proportional to the frequency of minor alleles in the target population. As f3 did not detect recent admixture events with two closely related ingroup tips, further tests were used to investigate introgression using smaller genomic windows.

#### Absolute divergence, d_XY_

Nei’s measure of absolute divergence (*d*_XY_) (Nei 1987) was used to compare genetic similarity between populations using 50 kb genomic windows for both simulated and empirical datasets. In all cases, population wide comparisons of *d*_XY_ were performed between; i) two ingroup populations (I_1_ and I_2_), ii) ingroup 1 and the outgroup (I_1_ and O) and iii) ingroup 2 and the outgroup (I_2_ and O). Comparions returing values approaching zero indicate high levels of similarity and a recent allelic split time. Low *d*_*XY*_ values are expected between ingroup samples (I_1_ and I_2_), or in cases where introgression may be occurring between an ingroup and outgorup.

Absolute divergence in simulated populations was calculated for each 50 kb window (n=500), again for admixture occuring at *f* = 0.0, 0.05, 0.1, 0.2 and 0.3. The distribution of *d*_*XY*_ values obtained for each comparison were plotted as histograms normalised for density by rescaling such that the maxima of the distribution is 1 (**Figure 4, S2, S3**). Introgression either from I_2_ into O or from O into I_2_ produced a decrease in absolute divergence across the genome, providing a benchmark for comparisons with empirical data. Admixture in the direction O to I_2_ provided a much clearer genome wide signal than the reverse direction, (I_2_ to O) indicating it would be easier to detect introgression from *P. australiana* into Australian *P. xylostella* than the reverse.

Based on the whole genome phylogeny (**Figure 1**), we expected mean *d*_XY_ between *P. australiana* and Hawaiian *P. xylostella* to be slightly lower than Australian *P. xylostella*. The *d*_XY_ distribution of empirical 50 kb windowed data provided no support of widespread introgression (**Figure 4C**), as values comparing the *P. australiana* outgroup with either *P. xylsotella* from Hawaii (I_1_) or Australia (I_2_) did not deviate from their expected values (**Table 3**). This suggests concordance with the whole genome tree topology.

**Table 3.** Confidence intervals (95%) for *d*_*XY*_ comparisons of populations

|  | *Pxyl.* Hawaii vs<br>*Pxyl.* Australia | *Pxyl.* Hawaii vs<br>*Paus.* Australia | *Pxyl.* Australia vs<br>*Paus.* Australia |
| --- | --- | --- | --- |
| ACT 2014 | 0.02101 - 0.02122 | 0.03788 - 0.03812 | 0.03828 - 0.03852 |
| ACT 2015 | 0.02102 - 0.02123 | 0.03721 - 0.03744 | 0.03771 - 0.03795 |
| SA 2014 | 0.02087 - 0.02111 | 0.03898 - 0.03926 | 0.03926 - 0.03955 |

#### Phylogenetic tree-tip distance proportions

The f3-statistic and *d*_*XY*_ were both used to test for introgression within populations. Next, we used the tree-tip distance proportion to assess whether evidence for introgression could be detected between individual sample pairs. Simulated genomes with introgression from I_2_ into O, or O into I_2_ at the rates *f* = 0.0, 0.05, 0.1, 0.2 and 0.3 were divided into 50 kb sequence alignments, as described above. Further subdivision was then performed so each 50 kb window contained just four sequences; two ingroup 1, one ingroup 2 and one outgroup sequence. This was repeated 64 times for each 50 kb window, then maximum likelihood phylogenic reconstruction performed for each alignment. Based on the whole-genome topology, we expected the outgroup to be a similar distance from both ingroup 1 and ingroup 2, unless introgression had occurred and shortened the distance between samples.

A proportion of the branch distance between I_2_ and O (*d*_*a*_) and I_1_ and O (*d*_*b*_) was then calculated for each phylogeny using equation 1, normalising values within the range 0-1. Tree-tip distance proportions are presented as histograms to graph the distribution (*x axis*), and normalised density (*y axis*). Although introgression from I_2_ into O (**Figure 5A, S4**) and from O into I_2_ (**Figure 5B**) were both detected using distance proportions, patterns did vary based on the direction of admixture. Clearer signals of admixture were evident in the direction O to I_2_ (**Table S4**), as this made ingroup 2 less similar to ingroup 1. Given I_1_ and I_2_ recently split, admixture from I_2_ into O is also expected to make the outgroup more similar to I_1_, decreasing detectability. Simulating introgression over five equaly spaced events was effective at detecting admixture using tree-tip distance proportions (**Figure S5**).

**Fig. 5.**
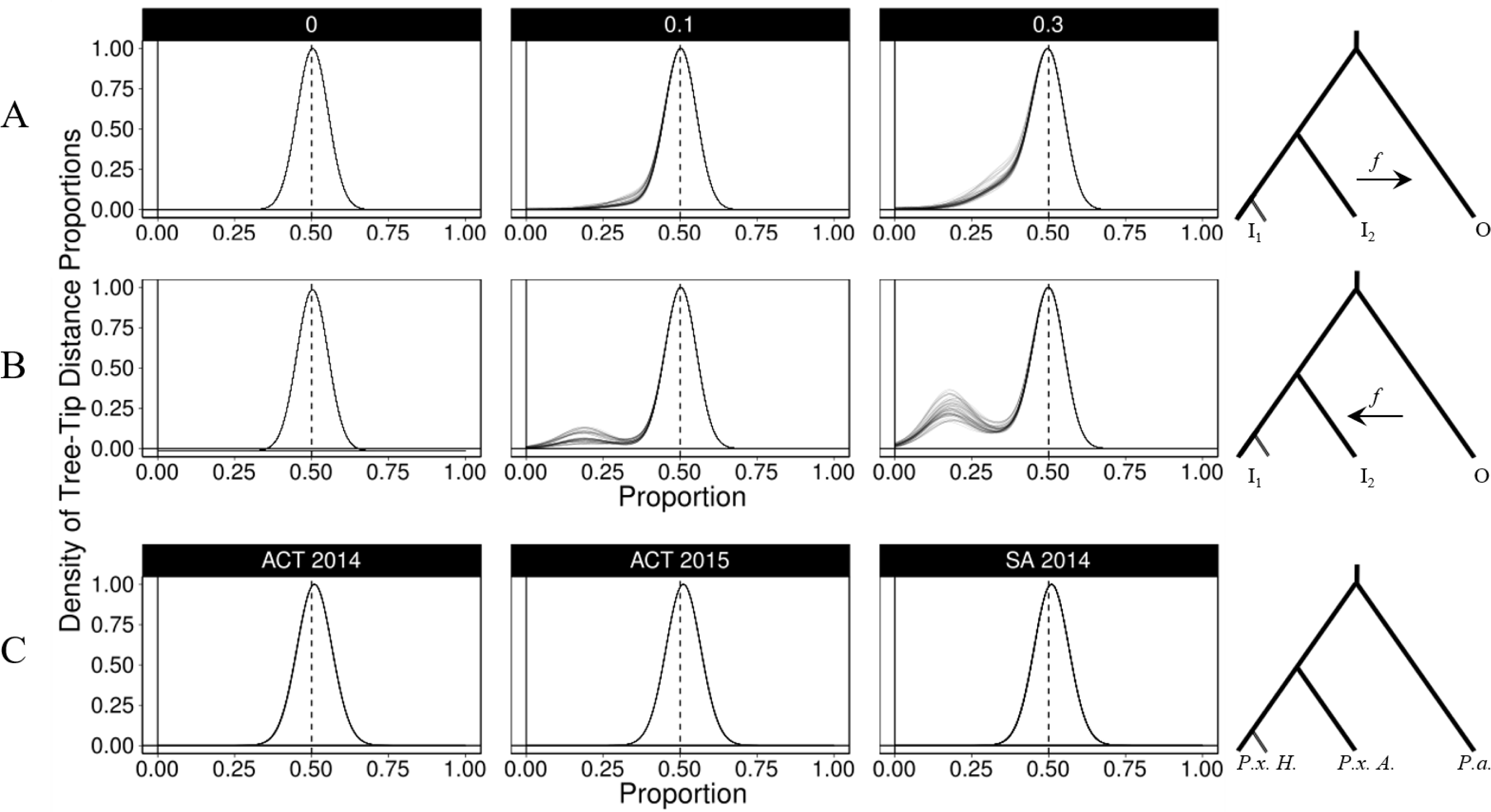
Histogram summaries of tree-tip distance proportions, depicting the phylogenetic distance between ingroup and outgroup sequences. A ratio of 0.5 indicates the outgroup sample has the same branch distance to both ingroup samples. Ratios close to zero indicate very short branch lengths between ingroup two and the outgroup, and are candidate regions for introgression. **A.** Simulated introgression from ingroup two to the outgroup at mixing frequencies of *f =* 0, 0.1 and 0.3. **B.** Simulated introgression from the outgroup into ingroup two at mixing frequencies of *f =* 0, 0.1 and 0.3. **C.** Empirical data for the three sympatric Australian *Plutella* populations (ACT 2014, ACT 2014, SA 2014). Each panel summarizes 7276 (±293) branch distance ratio calculations. The branch leading to ingoup 1 was standardized using the same two Hawaiian *P. xylostella* individuals for each of these comparisons.

This method was then applied to empirical data to identify genomic windows that were discordant with the species tree. Similar to the simulated datasets, genomic alignments were divided into non-overlapping 50 kb contiguous windows, then further subdivided into alignments of one *P. xylostella* and one *P. australiana* individual from sympatric Australian populations, plus two consistant *P. xylostella* from Hawaii. This produced 32 different sample combinations for each 50 kb window, including eight combinations from ACT 2014, eight from ACT 2015 and 16 from SA 2014. An average of 7276 (±293) maximum likelihood phylogenies were then produced for each of the 32 sample combinations to identify potential admixture that was not fixed in the population. A near-symetric and unimodal distribution of tree-tip distance proportions was observed in all cases, with the ranges of each density curve showing a large degree of overlap (**Figure 5C, Table S5**). Mean tree-tip distance proportions for each comparison were consistant within and between populations ACT 2014 (0.5088 - 0.5104), ACT 2015 (0.5102 - 0.5117) and SA 2014 (0.5086 - 0.5099). All proportion means were above 0.5 showing the Hawaiian *P. xylostella* had on average shorter branch lengths to *P. australiana*, consistant with the nuclear genome phylogeny (**Figure 1**). Under a widespread admixture hypothesis windows with distance prorortions below the mean (*P. australiana* closer to *P. xylostella* from Australia) should be much more frequent than above. The number of windows above and below three distances from the mean (0.05, 0.1, 0.15) was similar (**Table S6**) suggesting no clear evidence to support widespread admixture within *P. xylostella* and *P. australiana* individuals form three sympatric populations.

Despite lack of support for widespread hybridization and genome-wide introgression, we further investigated the tails of the tree-tip distance proportions at a distance 0.15 below the mean for each Australian population. These scaffolds (n=21) have the shortest branch lengths between *P. australiana* and sympatric Australian *P. xylostella*, relative to the branch length proprtions between *P. australiana* and Hawaiian *P. xylostella* (**Table S7**). Sliding window *d*_*XY*_ was performed on each of these scaffolds with 10 kb windows (sliding by 2kb), revealing just one region on scaffold KB207303.1 where Australian *P. xylostella* and *P. australiana* are more similar than between Hawaiian and Australian *P. xylostella*. Hostorical admixture between Australian *Plutella* species is one possible explanation for this result, although the region does not contain any protein coding genes (**Figure S6A**). A region on scaffold, KB207380.1, was identfied in the tree-tip distribition tail in 12/32 comparisons however *d*_*XY*_ indicated admixture across this region was unlikely (**Figure S6B**).

### Genomic regions with high interspecies divergence

The two *Plutella* species investigated in this study have been shown to have contrasting biologies and pest potential (Perry, et al. 2018), and although they can hybridize in laboratory crosses, we found no evidence for widespread admixutre among wild samples. This prompted us to ask which 50 kb genomic regions are most divergent between these species, and what kinds of genes do they encode? First, absolute divergence (*d*_XY_) between all *P. xylostella* and all *P. australiana* individuals was used to identify the top 1% most divergent genomic windows (**Table S8**). These included fifty-one 50 kb windows dispersed across 41 unique scaffolds and showed 33-61% greater absolute divergence than the genome-wide average (*d*_*XY*_=0.0369). Second, the top 1% of genomic windows showing highest divergence in nucleotide diversity (F_ST_) were also identified, showing values 70-110% higher than the mean (F_ST_=0.356). These two estimates of divergence only detected one 50 kb region common to both *d*_XY_ and F_ST_ (KB207411.1; 400,001..450,000 bp). Most windows with the highest *d*_*XY*_ had relatively low F_ST_ values, suggesting the two species share similar levels of polymorphism across these regions. Non-redundant protein coding genes (n=176) within these divergent genomic windows contained genes required for feeding including digestion (eg. chymotrypsin, trypsin, aminopeptidase-N), detoxification (eg. cytochrome P450s, carboxylesterases) and also gene regulation (zinc finger proteins) (**Table S9**). However, Tajima’s D showed most of these windows were within the genome wide 95% confidence intervals, indicating these regions are not likely to be under directional or balancing selection (**Figure S7, S8**).

## DISCUSSION

The discovery of cryptic species can often be inadvertent and arise from sequencing mitochondrial or nuclear amplicons (Landry and Hebert 2013; Stuart, et al. 2006), as well as whole genomes (Janzen, et al. 2017). The fortuitous identification of *Plutella australiana* was unexpected and raised initial concern over its pest status and whether specific management practices were required. *P. australiana* populations collected from across southern Australia (Perry, et al. 2018) and Australian *P. xylostella* (Endersby, et al. 2006) lack genetic structure, showing these species are highly mobile. Adaptive introgression of advantageous traits from one of these species into the other could potentially spread across the Australian continent. Despite high levels of movement, we sampled from sites where *Plutella* populations co-occur to attempt to detect either historical admixture or very recent hybridization.

The physical genome size of *P. xylostella* is estimated at 339 Mb (Baxter, et al. 2011) while the reference genome assembly is 393 Mb (You, et al. 2013) and includes sequencing gaps totaling ∼50 Mb. After aligning all resequenced genomes to the *P. xylostella* reference, only 170 Mb was retained in this analysis, which is likely to be caused in part by the sequence gaps and also high levels of genetic diversity (You, et al. 2013). Mapping *P. australiana* sequence reads to the *P. xylostella* genome is affected by mapping bias, as the most divergent loci will not map to this reference. This causes all branches to be shortend towards the reference, underestimating the diveregence between *P. australiana* and *P. xylostella* in the whole genome phylogeny. However, the introduction of this bias is unavoidable as the only reference geneome within the superfamily Yponomeutoidea is currenlty *Plutella xylostella.*

*Plutella australiana* were more similar to *P. xylostella* samples from Hawaii than Australia, based on shorter phylogenetic branch lengths for nuclear genomes and subsequent tree-tip distance proportions, lower F_ST_ values and lower f3-statistics. A better understanding of migration or transport routes enabling *P. xylostella* to colonize the world would help explain why this is the case. Several studies have found Australian *P. xylostella* mtDNA genomes have very low levels of diversity (Juric, et al. 2016; Perry, et al. 2018; Saw, et al. 2006), which is indicative of a population bottleneck (and other factors), while we found Hawaiian mtDNA genomes to be quite diverse. This suggests the Hawaiian Islands may have been colonized by a larger founding population, or multiple, independent invasions while Australia may have simply been colonized by a derived population of *P. xylostella* (Juric, et al. 2016).

Mitochondrial diversity between the two *Plutella* species was originally found to be ∼8.2%, based on sequencing COI amplicons (Landry and Hebert 2013), although the level of diversity across all thirteen protein coding genes is less (4.95%). This level of diversity was not sufficient to result in complete reproductive isolation between the two sister species when reared in the laboratory (Perry, et al. 2018). Using the 13 mitochondrial genes, we estimated the split time of *P. xylostella* and *P. australiana* to be ∼1.96 million years. To date, *P. australiana* has only been detected in Australia, yet this relatively recent split questions whether *P. australiana* did evolve within Australia. This would require a considerable migratory event some two million years ago from the ancestral *Plutella* source population to Australia, and no further migration. Future molecular screening of *P. xylostella* may identify cryptic *P. australiana* in other countries.

Phylogenies of genes or genomic windows can deviatie from an expected consensus topology or species tree and can be used to identify genomic regions that may be of biological interst. For example, genomic regions subject to incomplete lineage sorting (Scally, et al. 2012), horizontal gene transfer (Moran and Jarvik 2010) and adaptive introgression (Wallbank, et al. 2016) all produce discordant phylogenies. Despite simulated data detecteting minor levels of introgression using phylogenetic tree-tip distances across the genome, we found few discordant distances between individual *Plutella* samples across 50 kb genomic windows. The methods used here were not sufficient to reject small regions of decreased *d*_*XY*_ between Australian *Plutella*, which may be signals of past admixture. Future work into the evolutionary history of *Plutella* moths and sequencing outgroup genomes of *Plutella* species, will enable further analysis of these regions using the D statistic and *f*_*d*_ (Martin, et al. 2015).

The most divergent genomic windows between the two *Plutella* species identified using *d*_XY_ or F_ST_ showed little evidence for current selection and may potentially contain genes that underwent selection after speciation. These genes may reflect different abilities to evade host plant defenses or host plant preference, as many are involved with digestion and detoxification. Using absolute genetic divergence (*d*_XY_) to identify the most divergent genomic windows between *P. australiana* and *P. xylostella* may also be highlighting loci that are highly polymorphic or rapidly evolving. Further understanding of *Plutella* biology including mating timing, evolutionary history, host plant preference and behavious may provide further insight into these divergent loci.

*Plutella australiana* and *P. xylostella* are likely to have been in secondary contact in Australia for over 125 years (>1000 generations). Despite this, we found no support for widespread admixture, and although we can’t predict the amount of time these species have spent in geographic isolation, strong reproductive barriers are apparent in the field. Furthermore, *P. xylostella* and *P. australiana* will be a useful system to investigate the genetic basis of biological differences between cryptic species from an agricultural perspective.

## Acknowledgements

This work was supported by the Australian Research Council (grant numbers DP120100047, FT140101303) and Grains Research and Development Corporation (UOA1711-004RSX). CMW is supported by The Commonwealth Hill Trust and Grains Research and Development Corporation. Supercomputing resources were provided by the Phoenix HPC service at the University of Adelaide. We thank Kym Perry (University of Adelaide, Australia), Kevin Powis (South Australian Research and Development Institute, Australia) and Ron Mau (College of Tropical Agriculture and Human Resources, University of Hawaii) for sample collection along with Paul Hebert (Centre for Biodiversity Genomics) and Jean-François Landry (Canadian National Collection of Insects, Arachnids, and Nematodes) for the images of *P. australiana* and *P. xylostella* respectively. We also thank two anonymous reviewers for their comments on a previous version of this manuscript.

